# Genome assembly, transcriptome and SNP database for chum salmon (*Oncorhynchus keta*)

**DOI:** 10.1101/2021.12.27.474290

**Authors:** Eric B. Rondeau, Kris A. Christensen, Dionne Sakhrani, Carlo A. Biagi, Mike Wetklo, Hollie A. Johnson, Cody A. Despins, Rosalind A. Leggatt, David R. Minkley, Ruth E. Withler, Terry D. Beacham, Ben F. Koop, Robert H. Devlin

## Abstract

Chum salmon (*Oncorhynchus keta*) is the species with the widest geographic range of the anadromous Pacific salmonids,. Chum salmon is the second largest of the Pacific salmon, behind Chinook salmon, and considered the most plentiful Pacific salmon by overall biomass. This species is of significant commercial and economic importance: on average the commercial chum salmon fishery has the second highest processed value of the Pacific salmon within British Columbia. The aim of this work was to establish genomic baseline resources for this species. Our first step to accomplish this goal was to generate a chum salmon reference genome assembly from a doubled-haploid chum salmon. Gene annotation of this genome was facilitated by an extensive RNA-seq database we were able to create from multiple tissues. Range-wide resequencing of chum salmon genomes allowed us to categorize genome-wide geographic variation, which in turn reinforced the idea that genetic differentiation was best described on a regional, rather than at a stock-specific, level. Within British Columbia, chum salmon regional groupings were described at the conservation unit (CU) level, and there may be substructure within particular CUs. Genome wide associations of phenotypic sex to SNP genetic markers identified two clear peaks, a very strong peak on Linkage Group 15, and another on Linkage Group 3. With these new resources, we were better able to characterize the sex-determining region and gain further insights into sex determination in chum salmon and the general biology of this species.

## Background

Pacific salmon of the genus *Oncorhynchus* are iconic, culturally important keystone species spawning across freshwater watersheds that feed the Northern Pacific Ocean. Predominately anadromous, members of most species spend years at sea, consuming marine nutrients that are eventually deposited into coastal ecosystems where they provide a valuable source of food to numerous marine and terrestrial species as the salmon spawn and then die [1].

Chum salmon (*Oncorhynchus keta*) are the second largest of the Pacific salmonids and may have historically represented up to 50% of the salmonid biomass in the Pacific Ocean [2]. It is the most widely distributed of the Pacific salmonid species [3, 4], with spawning grounds ranging from Japan and the eastern coast of the Korean Peninsula through to Northern Russia, and from the Mackenzie River south through Central California in North America [5]. Among the most significant species of Pacific salmon in commercial fisheries – in an analysis of British Columbian commercial fisheries 2012-2015, chum salmon was the most plentiful species by weight in 3 out of 4 years analyzed, and second most valuable by processed value when averaged across the four year period ($31 million per year) [6].

A key and fascinating biological feature in salmonids is homing, whereby adults demonstrate an ability to return to the same riverine sites where they were spawned, although not all species show the same degree of site fidelity (reviewed in [7]). Some species, such as Sockeye, have been observed to return to within metres of where they were hatched (e.g., [8]), but other species vary in their fidelity to site of return and stray rate. Reasons for straying are likely varied (reviewed in [7]), but significant factors are thought to be juvenile freshwater residence time and freshwater migration distance, both of which lead to reduced imprinting. With chum salmon having relatively short freshwater residence (they migrate to sea as fry) and short migration distances (on average), it is perhaps not surprising that chum tend to have higher than average stray rates among the Pacific salmonids [7]. The consequences of such straying are that while regional-level differentiation (e.g., [9, 10]) and run-timing differentiation between summer and fall runs (e.g., [11–13]) can be observed, population-level genetic differentiation is not often seen within chum salmon.

The genomes of salmonids, including chum salmon, possess a key feature shared by all salmonid genomes, a salmon-lineage specific whole-genome duplications (WGD). WGDs very likely play one of the more significant roles in evolutionary innovation [14–17] and are found in plants (reviewed in [18]), fungi [19, 20], arthropods [21, 22], basal vertebrates ~500 million years ago (mya) [15, 23, 24], fishes ~300 mya [25–27], and more recently in ancestral salmonids ~90 mya [28, 29]. These major genome expansions have been proposed to allow for adaptations to new niches or conditions, particularly in times of major environmental change (reviewed in [30]). The occurrence of over 70 different salmonid species lineages stemming from the relatively recent ancestral WGD [29] offers a valuable system to i) observe evolutionary consequences of a relatively recent autopolyploid WGD, ii) identify ensuing mechanisms for regaining stable meiosis and cell division by regaining a functional diploid state through re-diploidization, and iii) draw associations between mechanisms of re-diploidization to potential genetic specialization that allow for species adaptation such as disease resistance. Additionally, each species has evolved unique morphology, life history strategies, and responses to common salmon pathogens (e.g., varied resistance to salmon aquaculture from pathogens such as the sea louse [31, 32]). This phenotypic variety provides future opportunities for exploring the biology and genetics behind the genomic architecture of whole-genome duplication have shaped these unique species.

The presence of these duplications, however, can present major technological challenges to genome assembly, due to limited differentiation between duplicated portions of the genome. Salmonids offer additional hurdles in that a significant portion of the genome still remains in a tetraploid-like state [33, 34], and may show lineage-specific re-diploidization patterns [35], or chromosome architecture through species-specific fusions [36]. While many challenges remain, the technological barriers to assembly of salmonid genomes are beginning to fall, as evidenced by the relatively rapid recent release of salmonid genomes [37–44]. A fully-annotated reference chum salmon genome will enhance development of genomics-based technologies to improve the effectiveness of fisheries management of the wild chum salmon fishery. This has already been performed for other Pacific salmon species in British Columbia (e,g., [45, 46]), and a genome assembly for chum would provide the ability to adopt similar management tools based on emerging high-throughput sequencing technologies.

Genetic resources in chum salmon have, as in many other species, been in a state of transition as genetic tools have advanced and become more widespread. Early work on population genetic structure in chum salmon utilized allozymes [47, 48] and microsatellite markers [9, 49] and provided the first range-wide studies on genetic diversity [10]. Recently, genetic stock identification tools have been shifting from microsatellites to single-nucleotide polymorphisms (SNPs), providing increased accuracy of genetic discrimination with increasing marker numbers [50]. Early identification of SNPs in chum salmon [51–55] led to the development of a SNP panel for assessing genetic diversity and population structures in chum salmon [13]; development of expanded SNP panels for fisheries management continues to occur with increased marker density and improving genetic baselines allow for increased power ([56]; Beacham T.D. and Sutherland B.J.G, Personal Communication). Restriction-site Associated DNA sequencing (RADseq) has recently enabled a much more rapid throughput for SNP discovery [57, 58], and studies in chum have utilized this technique to enable researchers to develop linkage maps to explore regions of residual inheritance associated with the aforementioned genome duplication event [34]. This advance in technology has further allowed for the identification of extended patterns of linkage disequilibrium, demonstrating the power of increased marker density on the identification of genomic features of large effect [59]. Despite this significant effort, unlike in other *Oncorhynchus* species (e.g., Rainbow trout [60]; Chinook salmon [41], sockeye salmon [61]), neither a whole-genome catalog of SNP markers nor whole-genome resequencing data has been available as a resource for chum salmon to date. The development of such a resource will further allow genetic resources, such as SNP panels, to be placed in context relative to genes or other annotated genomic features.

In this work, we have sequenced and assembled the genome of a mitotic gynogen doubled haploid chum salmon to eliminate allelic variation but retain paralog differences. Extensive multi-tissue RNA-seq was generated to provide the base for annotation of the genome as well as a tissue-specific expression atlas for future comparative studies. Finally, whole-genome resequencing was performed across 59 individual chum salmon from a select distribution of the species’ range to catalogue genome-wide diversity in this species. The utility of the dataset is further demonstrated by the genetic association of the sex phenotype onto the expected chromosome in a narrow window of elevated linkage disequilibrium.

## Methods

### Data availability

All raw sequencing reads and the assembled genome described in this project have been submitted to NCBI under BioProject PRJNA556729. SNP variant sets described below are available through Dryad repository.

### Animal care and sample collection

All animals were reared in compliance with Canadian Council on Animal Care Guidelines, under oversight from the Fisheries and Oceans Canada Pacific Region Animal Care Committee (PRACC). Chum salmon for genome sequencing and assembly and for transcriptome assembly were from Chehalis River Hatchery parents and reared at Fisheries and Oceans Canada in West Vancouver. Chum salmon mitotic gynogen doubled haploids were produced following procedures described by [62]. Briefly, eggs were fertilized with UV-irradiated sperm and pressure shocked (10,000 psi for 5 minutes) in batches at 30 min intervals between 4 and 7 hours post-fertilization. One individual from the 7h pressure shock group (Oke142-1, NCBI BioSample: SAMN12367893; Supplementary Table 1) was confirmed to be homozygous for maternal alleles using a panel of 14 microsatellites [49], and was used for genome sequencing and assembly (see below). The individual was euthanized in a bath of 200 mg/L tricaine methanesulfonate (TMS) buffered in 400 mg/L sodium bicarbonate prior to first feeding stage, and stored in ethanol before DNA extraction and whole genome sequencing.

For transcriptomic data, control Chehalis River Hatchery chum salmon produced from the same parents as Oke142-1 but without UV milt treatment or pressure shock were grown in aerated fresh well water in 200–3700 L tanks and fed hourly as fry and to satiation 3 times daily as parr with stage-appropriate manufactured salmon feed (Skretting Canada Ltd.). At approximately 7 months post-ponding, a single selected chum female (86.9g with a 19.3cm fork length) was euthanized with TMS as above, then rapidly (< three min, PRACC management procedure 3.7) team dissected to harvest 18 tissues (see Supplementary Table 2) for RNA extraction, with an additional tissue (testes) sampled from an juvenile male. All tissues were stored in RNAlater at −20°C until extraction. RNA extractions were performed using the Qiagen RNeasy Mini Kit following the manufacturer’s protocol.

For individuals used in resequencing, samples were obtained primarily through non-lethal sampling of fin clips or operculum punches from Fisheries and Oceans Canada hatchery brood programs. Additional samples were obtained from archived tissue sets used for genetic stock ID baseline development to supplement the dataset. In total, 59 individuals were utilized in this assessment, with DNA obtained via Qiagen DNeasy Animal tissue kit’s following manufacturer’s protocol) or phenol/chloroform extractions (following Thermo Fisher Scientific’s protocol for genomic DNA preparation [63]. Tissue types, sex, collection dates and locations are summarized in Supplementary Table 3.

### Genome sequencing and Assembly

DNA was isolated from RNAlater or ethanol preserved tissues using a phenol/chloroform extraction as per Thermo Fisher Scientific’s protocol for genomic DNA preparation [63]. Extracted DNA was submitted for genome sequencing across multiple library types, using both Illumina and PacBio sequencing instruments (summarized in Supplementary Table 1: SRA chum Gynogen). Extracted DNA was submitted to the McGill University and Génome Québec Innovation Centre (now the Centre d’expertise et de services Génome Québec) for construction of overlapping (library size estimate = 497 base pairs (bp) and non-overlapping (library size estimate = 620bp) IDT dual-indexed Illumina Shotgun libraries. Each library was sequenced twice on an Illumina HiSeq2500 on RAPID mode PE250. Extracted DNA was also submitted to the McGill University and Génome Québec Innovation Centre for construction of a single library of 10X Chromium linked-reads. Following library construction, the library was sequenced across three lanes of Illumina HiSeqX PE150. Extracted DNA was also submitted to the National Research Council Plant Biotechnology Institute Genome Core for Illumina mate-pair library construction and sequencing. Mate-pair libraries targeting 2-3kb, 4-6kb and 7-12kb were constructed, and sequenced on a lane each of Illumina HiSeq2500 PE125. Finally, extracted DNA was submitted to McGill University and Génome Québec Innovation Centre for construction of a Pacific Biosciences SMRT library using a sheared large insert library type, and the MagBead OneCellPerWell v1 collection protocol. The library was ultimately sequenced across 16 total SMRT cells.

Assembly protocols followed successful strategies utilized for Northern Pike e.g., [40, 61, 64, 65]. See Supplementary Table 4 for specific parameters to assembly and trimming that were tested. Reads were first trimmed for quality, adapters and minimum length using Trimmomatic [66], and BBmap’s FilterByTile was utilized to remove poorly performing portions of the Illumina reads (https://jgi.doe.gov/data-and-tools/bbtools/bb-tools-user-guide/bbmap-guide/; [67]). Allpaths-LG v52488 [68] was utilized with overlapping Illumina overlapping PE250 and Illumina mate-pair libraries using a 3.0 TB memory node on the Compute Canada cluster Cedar. Non-overlapping libraries were also included in two assembly attempts, but ultimately exceeded the memory availability on the node in the MergeNeighbourhoods2 module and were dropped in successful assemblies. Assembly parameters were primarily adjusted for coverage of each of the library types as had been performed in other species; additional modifications were made to read filtering to improve the assemblies.

Following Allpaths-LG assembly, scaffolds were passed into PB Jelly 2 v 15.8.24 [69] along with all subreads produced in PacBio sequencing. Nodes on Compute Canada’s Cedar cluster were used for all stages, with on-node temp directory and 48 cores used in all steps where allowed. Blasr parameters were ‘-minMatch 8 -sdpTupleSize 8 -minPctIdentity 75 -bestn 1 -nCandidates 10 -nproc 48 -maxScore −500 –noSplitSubreads’. Extraction.py was modified to ‘MAXGAPHOLD= 1000000’ to take advantage of memory available. Collection.py was run with ‘-m 3’. All other parameters remained default. Finally, the assembly was polished with Pilon [70] using the trimmed paired-end data, aligned to the genome utilizing ‘bwa mem –M’ and default parameters.

Scaffolds were ordered and oriented into chromosome representations (i.e., Pseudomolecules) predominately following the methods described in Christensen et al. (2018) [40]. The sequences underlying the markers for the published chum linkage map from Waples et al. (2016) [34] were aligned to the scaffold assembly utilizing BLAST (-outfmt 6, -word_size 48, perc_identity 94, -max_hsps 100, -max_target_seqs 10 -evalue 1E-16). All scaffolds with a link to at least one marker on the map were retained for subsequent pseudomolecule inclusion. Scaffolds were ordered and oriented to the extent allowed by the linkage map, although regions of low recombination limited the effectiveness of the maps alone at this task. Therefore, the sequences underlying the markers for the linkage map were also aligned to a higher contiguity genome of a related species (coho; GCF_002021735.2), and ordering and orientation was further refined based on the conserved synteny between the two species via manual review. Where discrepancies were observed, the chum linkage map was taken as correct to ensure major species-specific rearrangements were captured. Finally, pseudomolecules were aligned to genomes of additional salmonids rainbow trout GCF_002163495.1 [39], Atlantic salmon (GCF_000233375.1) [38], Chinook salmon (GCF_002872995.1) [40] and the non-duplicated outgroup to the salmonids, northern pike (GCF_000721915.3) [71] using Symap v4.2 [72] to ensure linearity was generally conserved, and where it was not, was supported by rearrangements observed in the linkage map.

A BUSCO v4.0.2 [73] analysis utilizing the actinopterygii_odb10 dataset and ‘-m geno –c 10 –sp zebrafish’ was used to analyze the gene representation within the assembly utilizing the RefSeq maintained assembly: GCF_012931545.1.

### Gene Annotation

Raw reads for RNA-seq libraries were uploaded into NCBI under BioProject PRJNA556729 for inclusion in the Eukaryotic Genome Annotation pipeline. NEBNext dual-indexed mRNA stranded libraries were constructed from tissues described above by the McGill University and Génome Québec Innovation Centre, and sequenced on a half lane of NovaSeq 6000 S4 PE150 (additional libraries in the lane consisted primarily of RNA-seq of Pink and Chinook salmon from related projects). Sequences were uploaded under: SRP216443, with individual accessions: SRR9841162 (Adipose), SRR9841163 (Brain), SRR9841160 (Gill), SRR9841161 (Head Kidney), SRR9841166 (Heart), SRR9841167 (Hindgut), SRR9841164 (Left Eye), SRR9841165 (Liver), SRR9841168 (Lower Jaw), SRR9841169 (Midgut), SRR9841171 (Ovary), SRR9841172 (Pituitary), SRR9841170 (Pyloric Caeca), SRR9841174 (Red Muscle Skin), SRR9841176 (Spleen), SRR9841177 (Stomach), SRR9841173 (Testes), SRR9841175 (Upper Jaw Nares), and SRR9841178 (White Muscle).

### Variant Calling

All individuals sequenced for variant calling (Supplementary Table 3) used Shotgun PCR Free IDT dual-indexed Illumina libraries, produced on a quarter lane of Illumina HiSeqX – library construction and sequencing were performed at the McGill University and Génome Québec Innovation Centre. Raw reads were uploaded to the NCBI BioProject PRJNA556729, with individual accessions listed in the supplementary table.

Variant calling followed the best practices pipeline of GATK 3.8 [74–76], and generally followed the methods previously outlined in Christensen et al. (2020) [61]. Raw paired-end reads were aligned to the scaffold-version of the genome (pre-pseudomolecule construction) using bwa (v0.7.17) mem [77] and the ‘-M’ option. Samtools (v1.9) [78] was used to sort and index the alignment files, while Picard (v2.18.9) [79] was utilized with the MarkDuplicates option to identify likely PCR duplicates, and with ReplaceSamHeader to add read group information to the alignment files. GATK’s HaplotypeCaller was then used ‘--genotyping_mode DISCOVERY –emitRefConfidence GVCF’ to generate gvcf files, GenotypeGVCFs was used to generate vcf on intervals, and CatVariants was used to concatenate interval files into a single vcf. A training variant set was generated using a hard-filtered subset of the first round of genotyping, utilizing VariantFiltration and the parameters ‘ --filterExpression “QD < 2.0 || FS > 60.0 | | MQ < 40.0 || MQRankSum < −12.5 || ReadPosRankSum < −8.0”‘ as well as the VCFtools (v0.1.14) [80] parameters ‘--maf0.1 –hwe0.01’. A truth set was generated by overlapping the linkage map SNPs with the hard-filtered training set to obtain SNPs found in both methods.

VariantRecalibrator was applied using these sets ‘-mode SNP -an QD -an MQ -an MQRankSum - an ReadPosRankSum -an FS -an SOR -an InbreedingCoeff’, and ApplyRecalibration run to generate a final SNP set ‘ --ts_filter_level 99.0’. Finally, as SNP calling began pre-pseudomolecule construction, vcfChromTransfer in the Genomics General repository (https://github.com/simonhmartin/genomics_general; commit: 9d12505) was used to lift over the VCF file based on the NCBI submission AGP file. This lifted-over VCF is included in the accompanying dataset as the “raw SNP” set (referred to as set 1 below).

VCFtools v1.14 [80] ‘--maf 0.05 --max-alleles 2 -- min-alleles 2 --max-missing 0.9 – remove-filtered-all --remove-indels’ was used to retain only bi-allelic markers with little missing data and remove the rarest variants (referred to as set 2). The next filter utilized the VCF.Filter.v1.0.py script [61] to remove variants with allelic imbalance ‘-ab 0.2’, followed by VCFtools to select only the 37 pseudomolecules ‘--chr’ (referred to as set 3). The final filter utilized BCFtools v.1.9 to filter variants for LD in a 20kb window ‘+prune, -w 20kb, -l 0.4, -n 2’ (referred to as set 4). Finally, VCFtools ‘–relatedness2’ was run to detect closely related individuals. In light of the results, individuals ‘ Oke180104-Fert164’ and ‘ Oke171107-D’ are recommended to be used cautiously in further analysis as they were deemed most likely to be haploid progeny (expected) and sibling (unexpected) respectively of other individuals in the analysis (can be applied to all sets prior to further analysis using ‘vcftools –remove’; removed for Figure 3 below, not removed in Figure 2).

**Figure 1:**
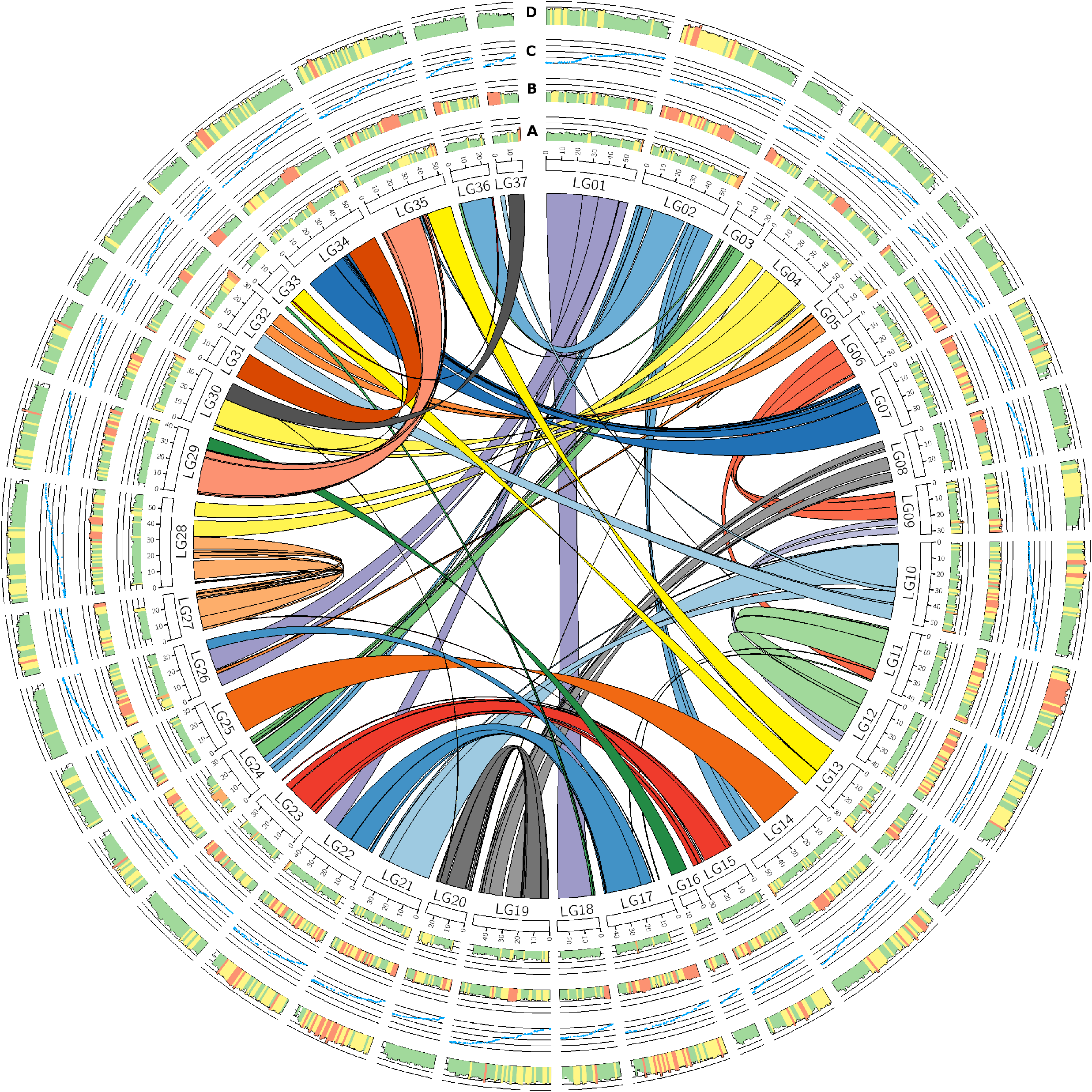
Circos plot of the chum salmon genome GCF_012931545.1. Inner ribbons demonstrate ohnologous regions (regions duplicated at the salmon-specific genome duplication event). Working in to out, Track A describes the average percent identity between the duplicated regions, in 1 Mbp bins. Track B describes the average percent identity in the chromosomes, in 1 Mbp bins. Track C describes the relationship to “Map 1” chum linkage map from Waples et al. (2016) [34]. Track D describes SNPs demonstrating elevated LD (R-squared >= 0.5) and >= 100 kb apart, demonstrated as a log10 based count, in 1Mbp bins.

**Figure 2:**
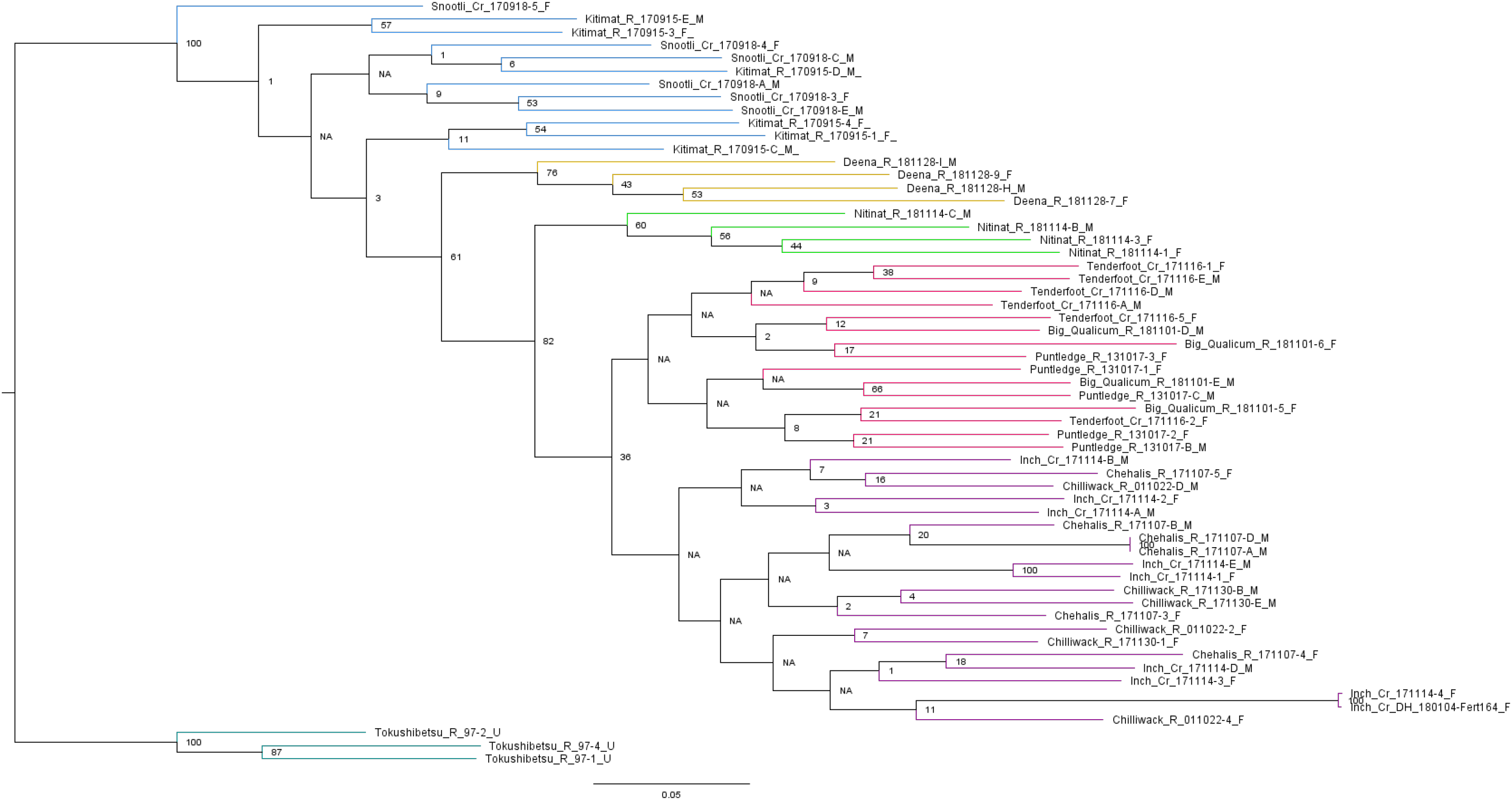
Dendrogram produced by SNPhylo, utilizing set 4 SNP data described in the text. Values at nodes indicate bootstrapping. Samples are coloured by geographic region.

**Figure 3:**
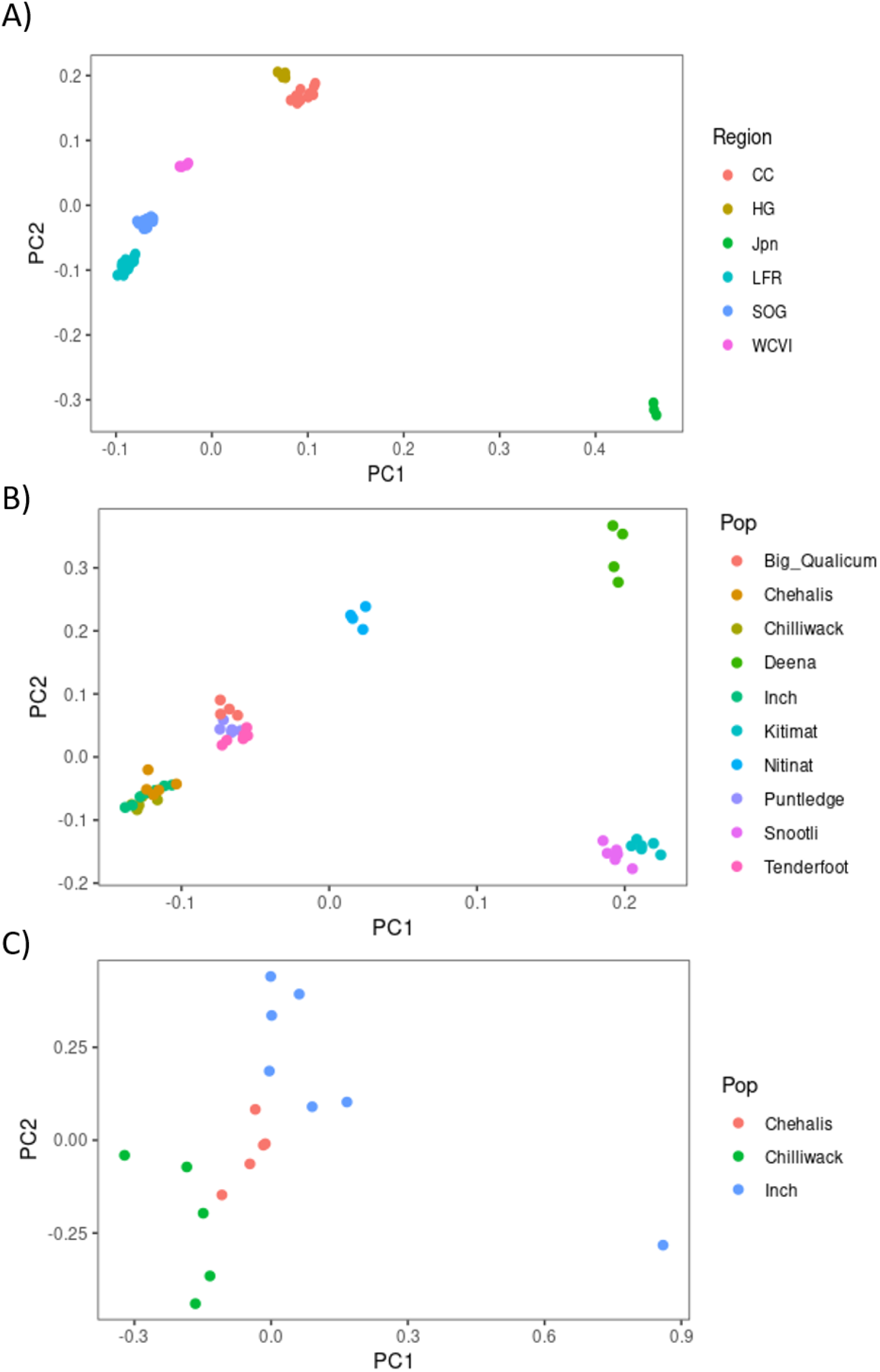
Principal component analyses performed on set 3 SNPs described in the text, using SNPrelate and plotted in ggplot2. Samples are coloured by collection and displayed in the legend. A) the full dataset (all samples) are presented. B), Japanese samples are removed from the analysis. C), the collections are reduced solely to the collections within the Fraser River drainage.

### SNP dataset analyses

SNPhylo [81] was run on the “set 4” dataset, using additional options ‘-m 0.05 -P Oket_chroms_37_ld0.2 -b –B 1000 -a 37’ in order to generate a bootstrapped phylogenetic tree of the chum salmon. Visualization was performed using the Figtree V1.4.4 package (http://tree.bio.ed.ac.uk/software/figtree/). PCA analyses were performed on the same dataset using the R package ‘SNPrelate’, with full and Canadian-only sample sets plotted–the set 3 is visualized in this work, with all visualization performed using the ggplot2 package [82].

The sex phenotypes associated with re-sequencing samples (Supplementary Table 3) were utilized as the basis for a genome-wide association analysis for sex. Utilizing the allele balanced SNP set (“set 2” above), VCFtools v1.14 was used to generate input for plink (chromosomes only). An association test was run in PLINK 1.9 [83] using the formatted output data, and resulting Manhattan plot visualized in R [84] using the qqman package [85]. Further visualization of identified SNPs were performed using the Adegenet package [86]. Counts of coverage utilized samtools v1.9 depth, using default parameters to calculate genome wide coverage over each individual *.bam alignment file, and using the ‘-b’ option to restrict the calculation to only the region of the growth hormone 2 gene (*GH2*) demonstrating elevated coverage in the males following a manual review of the alignments using IGV viewer 2.9.4 [87].

Duplicated regions, presumably from the Salmon specific 4R duplication event, were identified by alignments using the default settings of SyMap v4.2 [72], using a repeat-masked version of the genome following prior methods [61], by masking WindowMasker-based repetitive regions using ‘ sed -e ‘/^>/! s/[[:lower:]]/N/g’ from the RefSeq genome. Summary tracks were predominately generated using scripts from [40]: Orientation of the blocks were generated using Analyze_Symap_Block_Orientation.py; percent identity was determined using Analyze_Symap_Linear_Alignments.py; percent identify of repetitive regions identified using Percent_Repeat_Genome_Fasta.py. Linkage map markers from “Map 1” in [34] were aligned to the genome as previously described above using BLAST. Linkage disequilibrium (LD) was interpreted using the ‘--geno-r2’ option in VCFtools [80], and outputting only for those comparisons exceeding ‘--min-r2 0.5’ in order to identify the most highly linked SNPs – summaries were further limited to single chromosomes using the ‘--chr’ option. LD calculations utilized the allele balanced set (set 3) described above. LD track utilized counts of markers in linkage disequilibrium across at least 100kb, and summarized as a log sum per 1 million base pairs. Circos v0.69.9 [88] was utilized to visualize the data tracks described.

Heterozygosity analyses followed the same parameters and method as in [61]. Runs of homozygosity were identified from the variants that had been filtered for allele balance using PLINK v1.9 (parameters:—homozyg) [83]. The number of heterozygous genotypes and alternative homozygous genotypes per individual were counted using the same custom script described in the supplementary data of the sockeye genome [61]. Heterozygotes per kbp was calculated as the number of heterozygous genotypes divided by the total nucleotides in the genome (1,853,104,330) multiplied by 1 kbp. The heterozygosity ratio was calculated as the number of heterozygous genotypes divided by the number of alternative homozygous genotypes.

## Results and Discussion

### Genome Assembly and Annotation

From a raw data set consisting of 59X coverage (110 billion bp) of overlapping 250bp Illumina reads and 60X coverage (114 billion bp) of total mate-pair Illumina reads of three insert sizes (2, 5 and 8kb mean), multiple assembly attempts were performed varying the parameters on read depth as well as read-trimming. Ultimately, three of the attempts resulted in a completed assembly (see Table 1), with the final attempt being the most successful, with a contig N50 of 13.1 kb and a scaffold N50 of 653 kbp. Following AllPaths-LG assembly, contig gaps were filled utilizing PB Suite and 53 billion bp of Pacific Biosciences Sequel long-reads. Following Pilon polishing, utilizing the short insert Illumina libraries, scaffolds were organized into pseudomolecules representing the 37 chromosomes in chum salmon, predominately guided by the publicly available linkage map [34]; ultimately, the linkage map allowed for 70% of the genome assembly to be assigned to a linkage group, slightly lower but approximately equivalent to prior attempts in salmonids using equivalent techniques (e.g., [40, 61]). The final assembly was uploaded to NCBI under BioProject PRJNA556729 and ultimately was included in the RefSeq database as GCF_012931545.1.

**Table 1:** Assembly results for Allpaths and PBSuite based assemblies performed.

|  | Assembly size (Contigs) | Assembly size (Scaffolds) | Scaffold N50 | Contig N50 |
| --- | --- | --- | --- | --- |
| Allpaths | 1471097779 | 1813373414 | 653 | 13.1 |
| PBSuite | 1,766,907,823 | 1,852,809,593 | 665,581 | 52,191 |

Busco scores indicate that most of the genome is represented within the family, with results similar to what has been seen in Sockeye, with 85.0% complete (25.1% duplicated), 3.2% fragmented and 11.8% missing. This likely reflects the slightly more fragmented nature of the genome as compared to prior attempts using the same technology in other species. We believe this is most likely due to some minor shearing observed in the DNA utilized for library preparation. We did attempt to use 10X chromium data as part of this assembly process, but, our scaffolding power was negligible – after review, it is likely that DNA shearing noted in the bioanalyzer trace prior to library construction limited the size of the fragments from which to generate the linked reads, thus limiting scaffolding power. The raw data from this attempt is included under the BioProject (see Supplementary Table 1), but further attempts would need to use a separate individual in order to increase length of the starting material. Given that sequencing and assembly technology has advanced rapidly since we began this project, it is likely further efforts to improve the genome may benefit from the use of long-read technologies, where incredible advances in contiguity have already been demonstrated in salmonids [44, 89]. Indeed, a long-read assembly for chum salmon is planned by the authors, and will eventually replace this reference, in due course.

Following inclusion in the RefSeq database, the genome was annotated utilizing the NCBI Eukaryotic Annotation pipeline, ultimately yielding Annotation Release 100 (https://www.ncbi.nlm.nih.gov/genome/annotation_euk/Oncorhynchus_keta/100/) – see Table 2 for a summary. Gene annotation, via chum-specific reads, primarily utilized the 19 tissue RNA-seq dataset sequenced as part of this work (see Supplementary Table 2), with additional contribution of sequences from two additional datasets with publicly accessible RNA-seq data [90, 91]. Overall, gene numbers are comparable to other salmonid genomes, and thus likely reflect a relatively complete representation of the coding sequence. It is likely that a future genome that utilized long-reads would result in a slightly increased number of genes (as observed between *Oncorhynchus kisutch* Annotation releases 100 and 101, for example).

**Table 2:** Summary of Annotation Release 100 from the NCBI Eukaryotic Annotation pipeline. See https://www.ncbi.nlm.nih.gov/genome/annotation_euk/Oncorhynchus_keta/100/ for more details.

**Feature****Annotation Release 100**
|  |  |
| --- | --- |
| Genes and pseudogenes | 45643 |
| protein-coding | 36325 |
| non-coding | 6205 |
| transcribed pseudogenes | 222 |
| non-transcribed pseudogenes | 2821 |
| genes with variants | 14102 |
| immunoglobulin/T-cell receptor gene segments | 70 |

The resulting assembly is summarized visually in Figure 1. Duplicated regions, as identified via self-alignment using Mummer [92] reflect re-diploidized segments of the genome from the salmon-specific 4R duplication event. There are observations of elevated percent identity on the ends of some chromosomes (Figure 1) that demonstrate partial re-diplodization as in Waples et al. (2016) [34] (e.g., LG05 and LG32), but the effect is not nearly as extensive as that observed in other species. The repetitive elements identified by Window Masker were elevated in regions likely overlapping with centromeres based on synteny with other species for which chromosome arms have been described (Figure 1). Figure 1 also shows Map 1 from Waples et al. (2016) [34] (which can be further visualized in more detail in Supplementary Figure 1 and map 2 in Supplementary Figure 2), and demonstrates the co-linearity of the map with the pseudomolecules. As the maps do contain regions of low-recombination, much of the ordering and orientation of the scaffolds into pseudomolecules (but crucially, not the assignment to the pseudomolecule itself) relies heavily in some positions on the long-read and Hi-C based assembly of coho salmon (GCF_002021735.2). Given the extensive conserved synteny and co-linearity between orthologous salmonid chromosome arms demonstrated elsewhere (e.g., [36, 40]), this would appear to be a reasonable approach, and has been part of the development of pseudomolecules for short-read assemblies in salmonids previously (eg. [40]). Regions of the genome with high LD generally overlap with regions of reduced recombination as observed in the linkage map (Figure 1). Further exploration of regions of high LD can be observed in Supplementary Figures 3 and 4.

As a final clarification on the assembly presented, we note that pseudomolecules have been named within the publicly available assembly based on the linkage group naming mechanism in Waples et al. (2016), [34] to allow for direct comparison between the two works. However, the authors also note, and are enthusiastic about, the naming convention suggested by Sutherland et. al., [36] to describe chromosomal arms, and indeed the adoption of the system into the grayling genome assembly [42]. We provide here in Table 3 the naming for the pseudomolecules that could be suggested by such a system. While the pattern of fusions do make this system less than ideal, and the resulting chromosome names are somewhat unwieldy, we provide them here as a quick reference and potential guide to re-naming of the linkage groups should such a system continue to prove popular as future assemblies are released. Presenting both names here will hopefully ease future reference, whichever naming scheme ends up being formally adopted in future works.

**Table 3:** Pike-like chromosome naming for the chum salmon pseudomolecules described in this work, based on Sutherland et al. (2016) [36]

| **<u>Linkage Group</u>** | **<u>Accession</u>** | **<u>Alternate Naming</u>** |
| --- | --- | --- |
| Oket_LG01 | NC_050106 | 1.2-8.2 |
| Oket_LG02 | NC_050107 | 6.1-2.2 |
| Oket_LG03 | NC_050108 | 5.1 |
| Oket_LG04 | NC_050109 | 21.1-4.2 |
| Oket_LG05 | NC_050110 | 11.2 |
| Oket_LG06 | NC_050111 | 14.2 |
| Oket_LG07 | NC_050112 | 16.1 |
| Oket_LG08 | NC_050113 | 18.2 |
| Oket_LG09 | NC_050114 | 14.1 |
| Oket_LG10 | NC_050115 | 17.1-9.2 |
| Oket_LG11 | NC_050116 | 7.2 |
| Oket_LG12 | NC_050117 | 7.1 |
| Oket_LG13 | NC_050118 | 24.1-23.1 |
| Oket_LG14 | NC_050119 | 13.1-2.1 |
| Oket_LG15 | NC_050120 | 3.2 |
| Oket_LG16 | NC_050121 | 25.1 |
| Oket_LG17 | NC_050122 | 19.1 |
| Oket_LG18 | NC_050123 | 1.1 |
| Oket_LG19 | NC_050124 | 10.2-18.1 |
| Oket_LG20 | NC_050125 | 10.1 |
| Oket_LG21 | NC_050126 | 17.2 |
| Oket_LG22 | NC_050127 | 19.2 |
| Oket_LG23 | NC_050128 | 3.1 |
| Oket_LG24 | NC_050129 | 5.2 |
| Oket_LG25 | NC_050130 | 13.2 |
| Oket_LG26 | NC_050131 | 8.1 |
| Oket_LG27 | NC_050132 | 12.1 |
| Oket_LG28 | NC_050133 | 12.2-21.2 |
| Oket_LG29 | NC_050134 | 15.2-25.2 |
| Oket_LG30 | NC_050135 | 4.1-22.2 |
| Oket_LG31 | NC_050136 | 20.1 |
| Oket_LG32 | NC_050137 | 9.1-11.1 |
| Oket_LG33 | NC_050138 | 23.2 |
| Oket_LG34 | NC_050139 | 16.2-20.2 |
| Oket_LG35 | NC_050140 | 15.1-24.2 |
| Oket_LG36 | NC_050141 | 6.2 |
| Oket_LG37 | NC_050142 | 22.1 |

### Population level variation

Given the extensive distribution of chum salmon, attempts were made to maximize geographic distribution of the samples selected within the study. We were able to take advantage of an extensive collection of samples [10] in the archive of the Molecular Genetics Lab (Pacific Biological Station, Fisheries and Oceans Canada), combined with more recent contributions from various Fisheries and Oceans Canada hatchery staff for recent brood. While the collection is focused on British Columbia, the addition of the Japanese samples originating from the Tokushibetsu River on the Island of Hokkaido give a glance at the degree of variation expected across the Pacific. Samples and available metadata are summarized in Supplementary Table 3. In total, 15,372,999 nucleotide variants have been described with this data in the raw dataset, with described filters leaving 8,868,081 in set 2, 2,135,295 in set 3, and 94,080 in set 4. A summary of statistics by individual is given in supplementary table 5 [61]. On average (and ignoring the haploid individual), total lengths of runs-of-homozygosity (ROH) averaged 12.4 Mbp [0 - 40.8 Mbp as determined using default parameters]. Heterozygous SNPs per 1kbp averaged 1.47 (Standard Deviation = 0.15), while the heterozygosity ratio averaged 2.23 (Standard Deviation = 0.45). Overall, results are relatively similar to what was observed utilizing a parallel analysis in Sockeye salmon, although the overall length of ROH is lower (12.4 Mbp in chum salmon vs. 35.5 Mbp in Sockeye salmon), whereas heterozygous SNPs per 1kbp are increased (1.47 in chum salmon vs. 0.67 in Sockeye salmon), and the heterozygous ratio was approximately equivalent (2.23 in chum salmon vs. 2.21 in sockeye salmon after removing outliers). Deviations below the mean for both heterozygosity calculations were predominately associated with average coverage, implying that depth of sequencing likely impacted to some extent these calculations. Regardless, we demonstrate in chum salmon that there is a general increase in heterozygosity as compared to sockeye salmon, and establishes a comparative metric to be carried through to future comparative analyses in other Pacific salmonids.

Analyses of the SNP set resulting from whole genome resequencing (targeted coverage of 15X) should be considered exploratory, as collections were focused on geographic coverage to maximize variants within the catalogues rather than addressing additional questions. Nevertheless, the geographic variation explored allowed us to better understand differentiation among British Columbia locations. To this end, a bootstrapped maximum likelihood tree was constructed using a linkage-disequilibrium thinned SNP-set using SNPhylo [81]. As can be seen in Figure 2, the dendrogram clusters samples by regions similar to past analyses with more comprehensive sampling but using older marker technologies (see above). Samples can be resolved into regions corresponding to descriptions from the comprehensive sampling of Beacham et al. (2009) [10], with individual samples resolvable into Japan – Hokkaido; BC Central Coast (Snootli and Kitimat); BC-Haida Gwaii (Deena Creek); BC – West Coast Vancouver Island (Nitinat); BC – Strait of Georgia (Tenderfoot, Big Qualicum, Puntledge); and BC – Lower Fraser (Chilliwack, Inch, Chehalis). However, within clusters from multiple regions, we see a relative lack of resolution to the riverine level. Such observations are supported by Principal Component Analysis (PCA) as well (Figure 3); however, we do begin to see stronger delineation, possibly from the increased number of variant and dimensions in the PCA analysis. In Figure 3A, we observe differentiation across the Pacific Ocean (best described along PC1), and to a lesser degree geographically across the British Columbia coastline (along PC2). When described regionally, individuals from most populations can easily be resolved when focusing on the British Columbia coastline (Figure 3B), and we are able to see delineation among all collections, except those in the Fraser River Basin. Focusing on the Fraser River Basin sites alone, the pattern is less clustered (Figure 3C), although we do see some separation from salmon collected in different river systems of the Fraser River drainage.

Clustering techniques show that river-level resolution is not always observed. Such results have been noted in the past when considering fishery mixture resolution and describing assignments to region only (for example [9, 93]), but it is worth emphasizing that incomplete resolution among collected populations remains true when considering a relatively comprehensive genome-wide representation of variation. As part of the thinning procedure for SNPhylo, however, by default a significant number of SNPs are removed to increase the speed of the calculation. Alternatively, in the analysis of principal components, with just an LD threshold applied (0.2), a much greater number of SNPs were input into the resulting analysis, and it is likely that the number of SNPs in the end analysis played at least a partial role in the reduced delineation observed in the dendrogram relative to the PCA. While collection level differentiation does emerge in the PCA result, observations on reduced datasets (e.g., by chromosome) greatly inhibited the resolving power of the analysis (supplementary figure 5). Based on the results presented here, is is likely that collection level-specific SNPs could be identified in this dataset that maximize the population differentiation observed genome-wide, and that would further drive differentiation observed in the PCA. However, with such a small sampling size, it is likely that any such discovery would be more a representation of sampling depth, and the noise within a set would be high. However, this dataset is now available, should future researchers need to draw on a pool of potential SNPs from which to develop such assays.

Within BC, chum salmon regional groupings are described at the conservation unit (CU) level [94], and it is intriguing to note that there may be substructure to the results observed along those lines in the present analysis. For example, the Tenderfoot hatchery samples in the Howe Sound-Burrard Inlet CU do tend to cluster more strongly, relative to the other collection sites in the adjacent Georgia Strait CU suggesting that a greater sample size may allow recovery of further groupings. However, it is likely that straying, generally described as high in chum salmon, is playing a role in limiting genetic distinctiveness to the level of the CU (or higher) regional groupings. While sampling within the study focused primarily on large hatchery operations, it is also possible we are simply revealing a high degree of variation within each population due to a large effective population size, in which case sufficient additional sampling may coalesce around a mean per population. Still, even within the dataset here, the observation remains that individual population level resolution within a region may begin to be demonstrated with genome-wide representation.

### Mapping the sex-determining region

Although limited metadata was collected for individuals sampled beyond geographic locations sampled, we were able to collect phenotypic sex information on hatchery brood samples. Thus, we were able to explore genome wide associations (GWAs) of phenotypic sex. As demonstrated in Figure 4A, two clear peaks were observed with the GWAS: a very strong peak on Linkage Group 15, and another, albeit somewhat weaker, association on Linkage Group 3. As shown in Figure 4B, the specific region overlaps with an area of increased linkage disequilibrium on the distal end of LG15. In Figure 4C, the genotypes for each individual is displayed for the 20 SNPs seen as most associated with sex within the GWAS analysis. LG15 has been previously identified by McKinney et al. (2020), [59] as linked to sex during a RAD-seq based study of chum salmon populations within Alaska. In this prior work, linkage of sex to a particular region of the genome was complicated by two potential factors – a lack of a chromosome-level assembly for chum salmon, and the identification of a putative inversion along the chromosome that resulted in significant patterns of linkage. We utilized the sex-linked RAD loci to position the markers onto the new genome assembly and observed that while all were indeed placed along Oket_LG15, they appeared to be more dispersed along the chromosome, and were not strongly linked to sex within our geographically distinct sample set (Supplementary Table 7). Within the present study, we observed sex linked to a very narrow region along Oket_LG15; while some noise is observed, the peak is approximately in the 30.8 Mbp to 31 Mbp region and encompasses four annotated genes: potassium/sodium hyperpolarization-activated cyclic nucleotide-gated channel 2-like; E3 ubiquitin-protein ligase RNF126-like; SURP and G-patch domain-containing protein 1-like; and serine/threonine-protein kinase STK11-like. While we do not suggest any of these are the sex-determination gene – as with other Pacific salmonids it is presumed to be sdY [95] – given that the underlying genome assembly is female, this likely represents the approximate region where sdY is inserted on the Y-chromosome, and limited recombination surrounding the region has led to sex-specific markers extending to autosomal-like sequence flanking the insertion. This region (on chromosome 3.2 based on the naming scheme in Sutherland et al., 2016 [36] and Table 3) would appear to be a unique placement thus far in sdY mapping – however, the relatively common observation of sdY on chromosome arm 3.1 (sockeye salmon, coho salmon, lake whitefish; [96] and references therein) does suggest that inter-homeologue transfer between chromosome arms arising from the most recent salmon-specific duplication could be a mechanism for this transfer.

**Figure 4:**
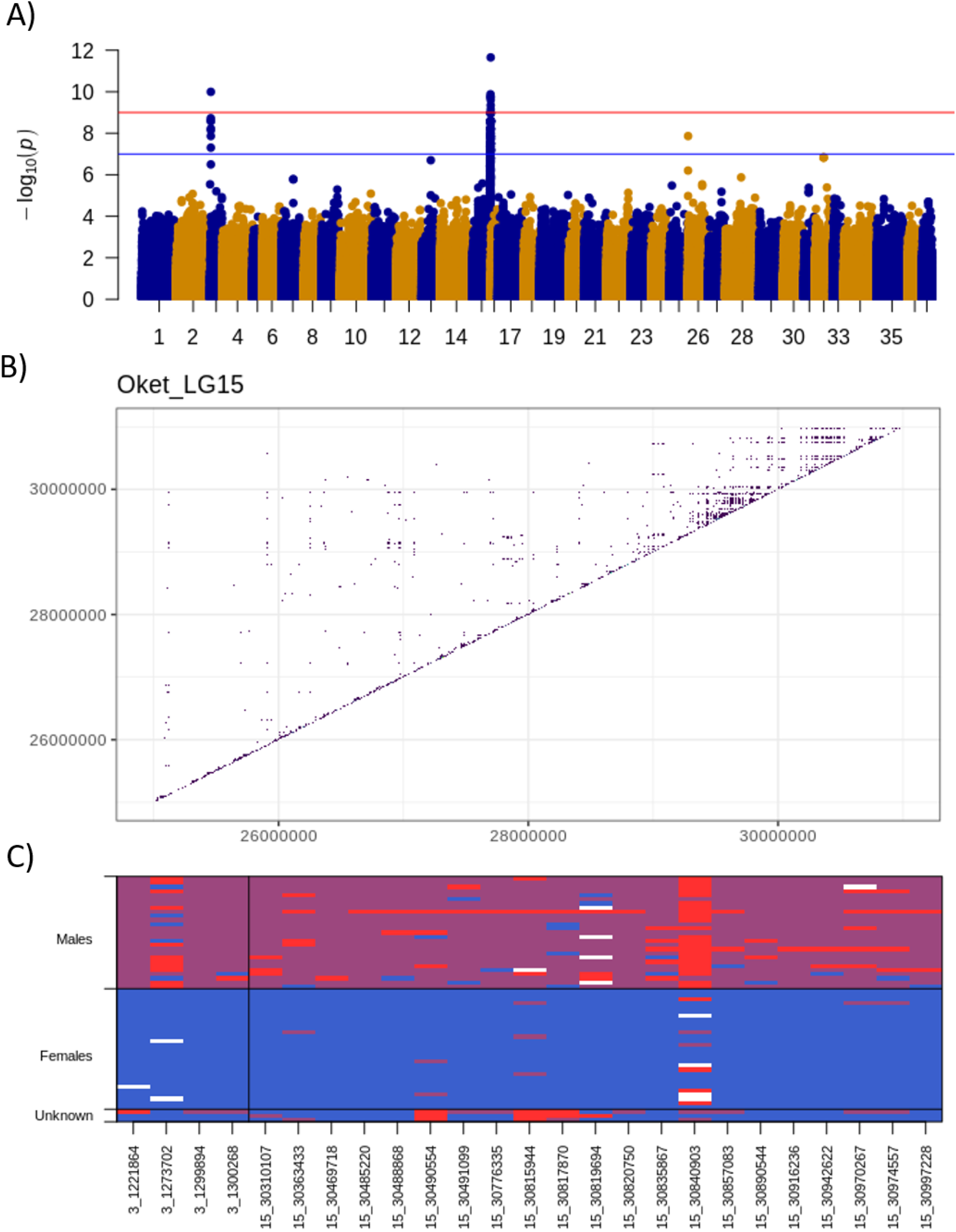
Association of the phenotypic sex to the genome utilizing SNP variant set 1. A) the results of the GWAS are presented, with Bonferroni-adjusted p-values shown at the 5% level (blue line) and 1% (orange line) levels. B) The SNPs with R-squared greater than 0.5 are counted, and plotted to show relationship of distance between SNPs being measured, for the region flanking the signal on Oket_LG15. C) The genotypes for each individual is displayed for the 20 SNPs seen as most associated within the GWAS analysis, with homozygous reference in blue, heterozygous in purple, homozygous alternate in red, and missing genotypes in white. Samples are sorted to group males, females and unknowns (Japanese samples—most likely females).

The strong secondary peak observed on Linkage group 3 is slightly more confounding and intriguing, as it does not appear to be linked to a known sex-determination orthologue in salmonids [96], and because potential sex-markers appear linked to those on LG-15. While it could be linked to a misplaced contig within the assembly, comparative mapping between additional species did not suggest anything was misplaced based on conserved synteny (data not shown; performed within Symap using default parameters) – if this is the case, it is likely that a future long-read based assembly will correct such a matter. It seems most likely in this case that it represents a repetitive element or otherwise duplicated sequence that is prominent in the Y-specific region but is not present in this female genome; thus, mis-mapping appears to occur elsewhere in the genome. A manual review of the region does imply a highly repetitive region, with great differentiation in depths indicative of collapsed repeats. Such mismapping based on collapsed repeats or a lack of sex-specific reference is not uncommon (e.g., as demonstrated in Chinook salmon by mapping of the Y-specific growth hormone pseudogene to the GH2 locus on a different chromosome [97]) and it may be that assembly of a male genome will reveal repeat patterns underlying this unexpected result observed here. There may be additional, more complex reasons based on the observance of multiple sdY regions seen in other species (e.g., Atlantic salmon [98]), although other explanations may be equally likely here. Observations have been made elsewhere that GH-Y, a commonly used proxy for genetic sex in salmonids [99], was found to be missing in males or present in females in some chum salmon populations [100]. While the presented genome is female-based (and thus not predicted to contain GH-Y), observation of relative coverage at the most closely related gene in the genome – GH2 – indeed implied that between 0-5 copies of GH-Y are observed in male individuals, with those males observed to be missing GH-Y being from Kitimat (2x), Snootli (1x) and Tenderfoot (1x): see supplementary table 6. These data do not suggest the phenotypes are mis-identified, however, as inclusion of a Rainbow trout sdY into the alignment phase demonstrated that the presence of sdY matches the phenotype, as would be predicted [95]. No copy-number differences could be interpreted from the sdY alignment unfortunately, as the underlying sequence from trout appeared too differentiated to obtain a reliable estimate of coverage; however, reads were observed aligned to the sequence in all male individuals and not in female individuals in a manual review utilizing IGV viewer. Still, the GH-Y results do indicate that there is variability in the genomic architecture surrounding sdY, and perhaps may indicate that alternate locations within the genome could be influential. Whatever the underlying genomic architecture of the sex-determination region may be in chum salmon, the result presented here underlines the usefulness and ease of use of the presented SNP dataset and reference genome in mapping a trait of interest to the appropriate chromosome and chromosomal region within the genome.

## Conclusions

The genome assembly for chum salmon represents a relatively complete representation of the chum salmon genome: the first such resource for the species. Contiguity and completeness is likely most affected in regions with high residual tetraploidy or incomplete re-diploidization. While long-read based assemblies (and future sequencing technologies) are likely to generate a more complete picture, the current genome assembly represents a valuable resource for chum salmon on par with those available for Chinook, sockeye, and longstanding assemblies for Atlantic salmon and rainbow trout that allowed a transformation in genomic understanding of these commercially and culturally specific species (e.g., [101]). Complementing the presented genome is a pilot-scale catalogue of variation that provides a genome-wide resource for British Columbian chum salmon populations, and allows for contrasting variation in Western and Eastern Pacific lineages. Such a dataset will be explored further as a resource for SNP genotyping panel expansion, structural variation discovery, or as demonstrated here, in identification of the chromosome and position most likely to contain the sex-determination gene in chum salmon.

## Acknowledgements

We would like to thank the staff at McGill University and Genome Quebec Innovation Centre (now the Centre d’expertise et de services Génome Québec; https://cesgq.com/) in Montreal, QC, Canada, and the NRC Plant Biotechnology Institute Sequencing Centre in Saskatoon, SK, Canada, for their work on library construction and sequencing on this project. Compute Canada (https://www.computecanada.ca/) provided much of the computing power for genome assembly and SNP discovery, primarily on the Cedar cluster. Support for this research from Fisheries and Oceans Canada, the Canadian Regulatory System for Biotechnology.

## Supplementary Data

Supplementary Table 1: Biosample and SRA data for individual chum used in generating the genome assembly.

Supplementary Table 2: Biosample and SRA data for individual chum used in generating the Illumina RNA-seq data.

Supplementary Table 3: Biosample and SRA data for individual chum used in generating the Re-sequencing data.

Supplementary Table 4: Allpaths-LG parameters explored in attempting to obtain the highest contiguity assemblies.

Supplementary Table 5: Heterozygosity metrics by individual are described. Includes counts of missing genotypes, Homozygous Reference and Alternate genotypes, Heterozygous genotypes, average depth at called sites, the mean count of heterozygous SNPs per kbp, the ratio of Heterozygous genotypes to Homozygous alternate, and the total length of runs-of homozygosity as determined from PLINK using default parameters.

Supplementary Table 6: Depth of coverage across the alignments, and at the GH2 locus to approximate the count of GH-Y copies in each individual. GH2 is used for this calculation due to the lack of GH-Y in the reference genome, and therefore the alignment of GH-Y to the closest homologue.

Supplementary Table 7: Placement of SNPs associated with phenotypic sex from McKinney et al. [59] in Alaskan chum populations onto the current reference genome.

Supplementary Figure 1: Plotting the association between Linkage groups in Waples et al. (2016), [34] map 1, and the reference genome assembly presented in this work.

Supplementary Figure 2: Plotting the association between Linkage groups in Waples et al. (2016), [34] map 2, and the reference genome assembly presented in this work.

Supplementary Figure 3: Plotting the linkage disequilibrium along each chromosome. SNPs are only displayed if R-squared is greater than 0.5, and is plotted as a count of SNPs.

Supplementary Figure 4: Plotting the linkage disequilibrium along each chromosome. SNPs are only displayed if R-squared is greater than 0.5, and each SNP is plotted by R-squared value.

Supplementary Figure 5: Principal component analyses performed on set 3 SNPs described in the text, using SNPrelate and plotted in ggplot2 and reduced to only query LG15. Samples are coloured by collection, displayed in the legend. In panel A, the full dataset (all samples) are presented. In panel B, Japanese samples are removed from the analysis. In panel C, the samples are coloured by collection site rather than by region.

